# Optimising risk-based surveillance for early detection of invasive plant pathogens

**DOI:** 10.1101/834705

**Authors:** Alexander J. Mastin, Timothy R. Gottwald, Frank van den Bosch, Nik J. Cunniffe, Stephen R. Parnell

## Abstract

Emerging infectious diseases of plants continue to devastate ecosystems and livelihoods worldwide. Effective management requires surveillance to detect epidemics at an early stage. However, despite the increasing use of risk-based surveillance programs in plant health, it remains unclear how best to target surveillance resources to achieve this. We combine a spatially explicit model of pathogen entry and spread with a statistical model of detection and use a stochastic optimisation routine to identify which arrangement of surveillance sites maximises the probability of detecting an invading epidemic. Our approach reveals that it is not always optimal to target the highest risk sites, and that the optimal strategy differs depending on, not only patterns of pathogen entry and spread, but also the choice of detection method. We use the example of the economically important arboreal disease huanglongbing to demonstrate how our approach outperforms conventional methods of targeted surveillance.

## Introduction

The collapse of the American chestnut population in the eastern United States in the early 20th century^1^; the English elm in 1960s and 1970s UK^2^; tan oak and coast live oak in the western United States over the last 30 years^3^; and the citrus industry in Florida since 2005^4^, have all resulted from the emergence of pathogens which were not previously present. Emerging infectious diseases (EIDs), such as these, are an increasing threat to wild and cultivated plants worldwide^5–7^. In some cases, EIDs may be established pathogens which have moved into new areas, as exemplified by the ongoing spread of the bacterium causing the citrus disease huanglongbing in the US^8^ and a competent vector of this pathogen in Europe^9^. In other cases, completely new pathogen strains may emerge, as seen with the emergence of the CoDiRO strain of *Xylella fastidiosa* in Italy^10^ and new strains of wheat stem rust worldwide^11^. Although changes in farming practices, land use, and climate, are important drivers of these processes^6^, most attention to date has focused on the global movement of people, plants, and plant products through travel and trade^12,13^. However, efforts to target these pathways directly through movement restrictions, quarantine, and border inspection^14^ often fail^15^, allowing pathogens to establish and spread beyond the threshold at which attempts to control the pathogen become infeasible^16^. As a result, it is increasingly recognised that surveillance activities must already be in place in the region of interest before pathogen entry^17^, and must be capable of detection before the prevalence (i.e. the proportion of the host population which is infected) exceeds the point at which control is almost certain to be no longer possible. This ‘maximum acceptable prevalence’ will be impacted by factors such as the epidemiology of the pest, its likely impact, and the availability and feasibility of control measures.

Developing an effective early detection surveillance strategy is complicated by the need to survey large, heterogeneous, areas of landscape over an indefinite timespan, in the face of limited financial and logistical resources. Whilst it is well accepted that attention must be focused on sampling enough hosts regularly enough for the prevalence at first detection to be likely to be acceptably low^18^, consideration must also be given to which hosts should be inspected and where to sample. One way of achieving this is through ‘risk-based’ or ‘targeted’ surveillance ^19,20^, in which the types of hosts or locations judged most likely to contain the pest or pathogen are preferentially selected for inspection or sampling^21^. Although the merits of targeted surveillance are well recognised, most work to date has focused on identifying static ‘high risk’ groups and locations using statistical models^22,23^. Although these methods of planning targeted surveillance are a valuable and versatile method of quantifying the infection risk amongst different groups, they do not explicitly account for the epidemiological processes which determine where and, importantly, when a pathogen will be present. As a result, there is a risk that the surveillance strategy may not be optimally targeted; resulting in low performance and/or excessive costs - both of which can ultimately lead to surveillance system failure.

The inability of conventional risk-based strategies to account explicitly for the epidemiology of the pathogen has important implications for early detection surveillance planning. To take a simple example, a common targeted surveillance strategy for EIDs is to focus on areas where the pathogen is more likely to first enter^24^. However, conventional methods do not tell us whether resources should all be placed around the single highest risk site, or spread across other potential introduction sites as well. Such questions can be answered by considering the placement of surveillance resources as an optimisation problem. By linking spatial and/or temporal simulation models which replicate the spread of the pest or pathogen to computational optimisation routines to identify particular sampling patterns, precise surveillance and/or control strategies which minimise the impact of the pest or pathogen can be identified^25^. Whilst much work to date has focused on identifying how best to conduct surveys in order to achieve certain disease management or mitigation objectives^26–30^, there has been little work on how to spatially target surveillance resources in order to detect pathogens at an early stage. No previous study has addressed the pivotal question of: where should surveillance resources be located to maximize the probability to detect an invading pathogen before it reaches a certain prevalence?

We propose and test a novel approach to surveillance planning which explicitly accounts for the spatial spread of a pathogen. Using a spatially-explicit, stochastic, epidemiological model, we represent the processes of pathogen introduction and onward spread across a real-world host landscape, continuing these simulations until a pre-defined threshold prevalence is exceeded. We then couple these simulation outputs to a stochastic optimisation algorithm designed to select those surveillance sites which maximises the probability of detection of the pathogen, allowing for a range of logistical parameters, such as different sampling intensities and detection abilities. We use this optimisation method approach to answer the question of where in a landscape surveillance should be targeted, focusing on the following questions:

i. How easy is it to implement this new strategy, and how much of an increase in detection probability can it achieve, compared to conventional site selection approaches?
ii. How do the locations and frequency of pathogen introductions influence the optimal arrangement of surveillance locations?
iii. What is the impact of the number of survey sites, the frequency of surveys, and the diagnostic sensitivity of the detection method on the optimal arrangement of surveillance locations?
iv. Can we identify general surveillance designs that approximate the optimised surveillance schemes and thus could be readily deployed in practice?

To demonstrate our method in the context of a pressing example, we use huanglongbing (syn. citrus greening, HLB) – a high profile, devastating disease of citrus trees – in the US State of Florida as a case study. HLB is caused by the bacterium *Candidatus* Liberibacter asiaticus (Las) and spread by the Asian citrus psyllid, *Diaphorina citri*, which has been established in Florida as an invasive species since at least 1998^31^. HLB is currently endemic in the state where it decimated the citrus industry in less than a decade, following first detection in 2005^32^. We consider here a scenario prior to this incursion, in which the psyllid is present but Las is absent from the state, in which there is an immediate risk of introduction of Las through human movements from other currently infected areas (such as Brazil and China). We use different estimates of where and how often the pathogen is introduced to the state in order to capture the inherent uncertainty in these processes and investigate how these influence the optimal surveillance strategy.

## Results

Although our spread model identifies a number of areas of high risk of pathogen presence (predominantly centered around areas of high citrus density – Figure 1), when we apply the optimisation routine we find that the common practice of focusing surveillance in a small number of highest risk areas generally does not maximise the mean probability of detecting the pathogen (Figure 2). For any combination of epidemiological and surveillance parameters, the conventional method of selecting sites based on site-specific ‘risk metrics’ gives a consistently lower probability of pathogen detection than the optimised approach (even over 1,000 realisations of these alternative sampling methods - see Figure 3). Moreover, we show that the effectiveness of the detection method used to find disease determines how that method should be deployed across the landscape; with poorer performing methods often requiring a greater focus in a relatively small number of risk ‘clusters’ (Figure 4). Our method is also robust to misspecification of model parameters, and consistently outperforms alternative methods in these situations (Figure 5). These results clearly demonstrate the importance of carefully considering pathogen epidemiology and entry processes, but also the detection efficiency of inspection and detection technologies, in a holistic manner when planning early detection surveillance.

**Figure 1.**
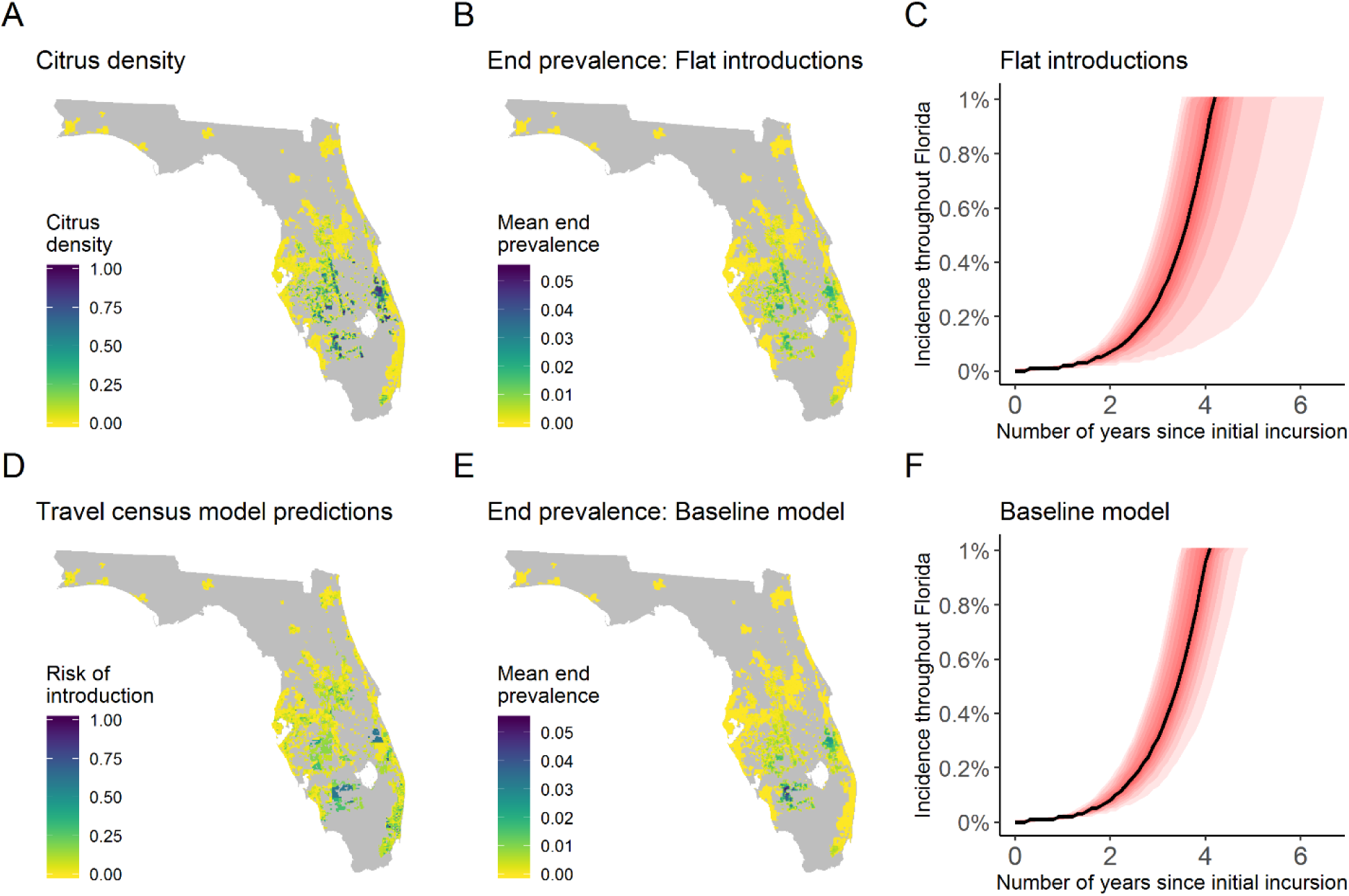
Simulation model upon which the optimisation is based. (A) Distribution of citrus trees (which also represents the relative rate of introduction under the assumption of a ‘flat’ distribution of pathogen entry). Plot D shows the relative risks of introduction according to the ‘travel census’ model (which is combined with the citrus density to estimate the relative distribution of introductions under the ‘variable’ model). Plots B and E show the mean end prevalence if introductions are based only on citrus density (‘flat’ pathogen entry, B) or both citrus density and travel census risk (‘variable’ pathogen entry, E). Plots C and F show the 5th-95th percentiles of the disease progression curves under each of these assumptions, with greater intensity of colouration for percentiles approaching the median (shown as a solid line).

**Figure 2.**
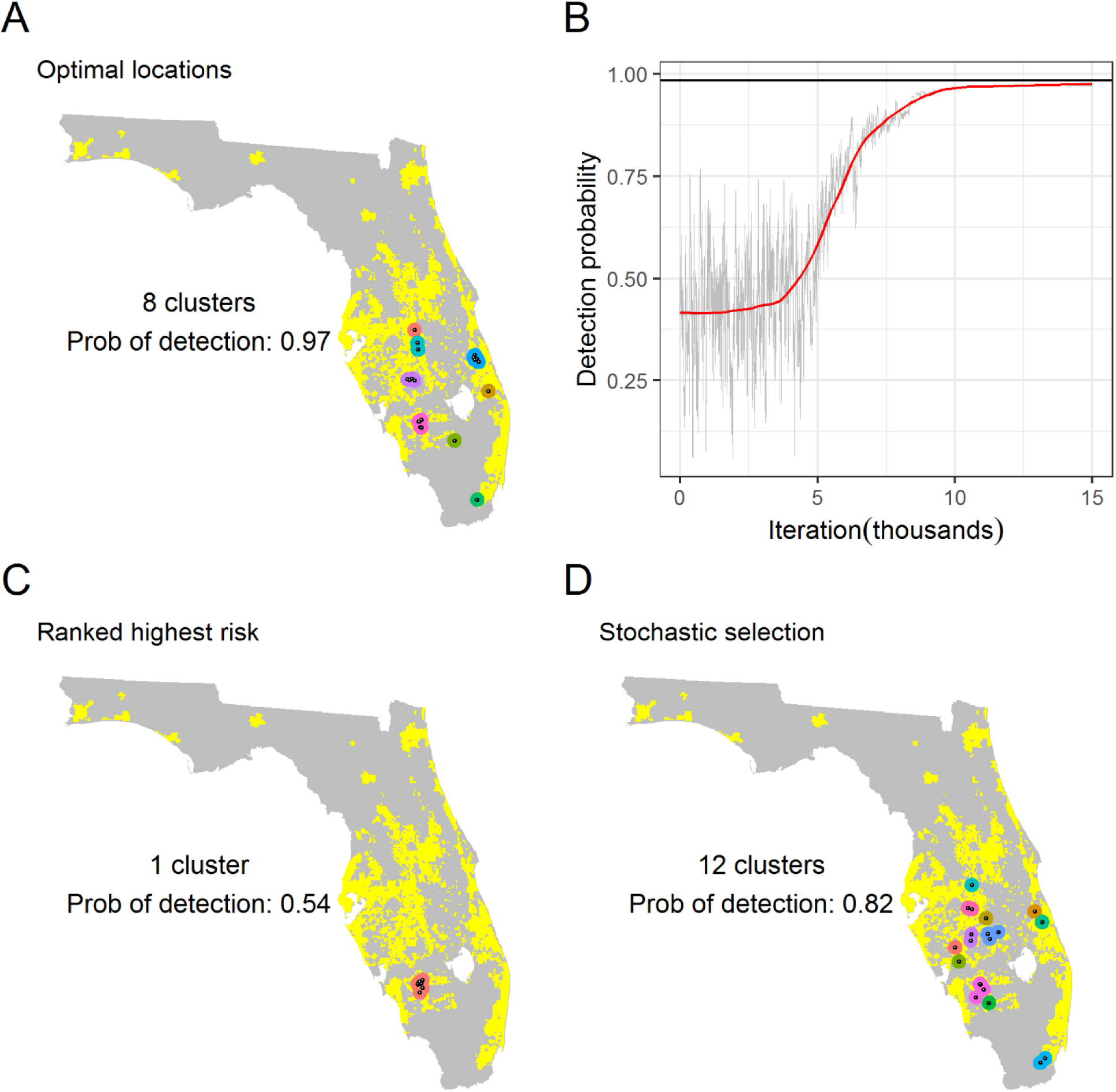
Arrangement of sampling sites under different selection schemes. These plots show the distribution of selected sites for one case of optimised selection, and two alternative risk-based approaches based upon the mean end prevalence obtained from the simulation model. Panel A shows the sites selected to maximise the probability of detection when using simulated annealing. The progression of the detection probability over the first 15,000 iterations of the simulated annealing algorithm is shown in panel B, with the solid black line indicating the final detection probability after 100,000 iterations. Panel C shows the 20 sites with the highest mean end prevalence over all realisations, and panel D shows one arrangement of 20 sites selected with a probability proportional to the mean end prevalence. Clusters (defined as sites within 20km of each other) are shown in plots A, C, and D. Estimates of the number of clusters and the probability of detection under the different sampling patterns are also shown.

**Figure 3.**
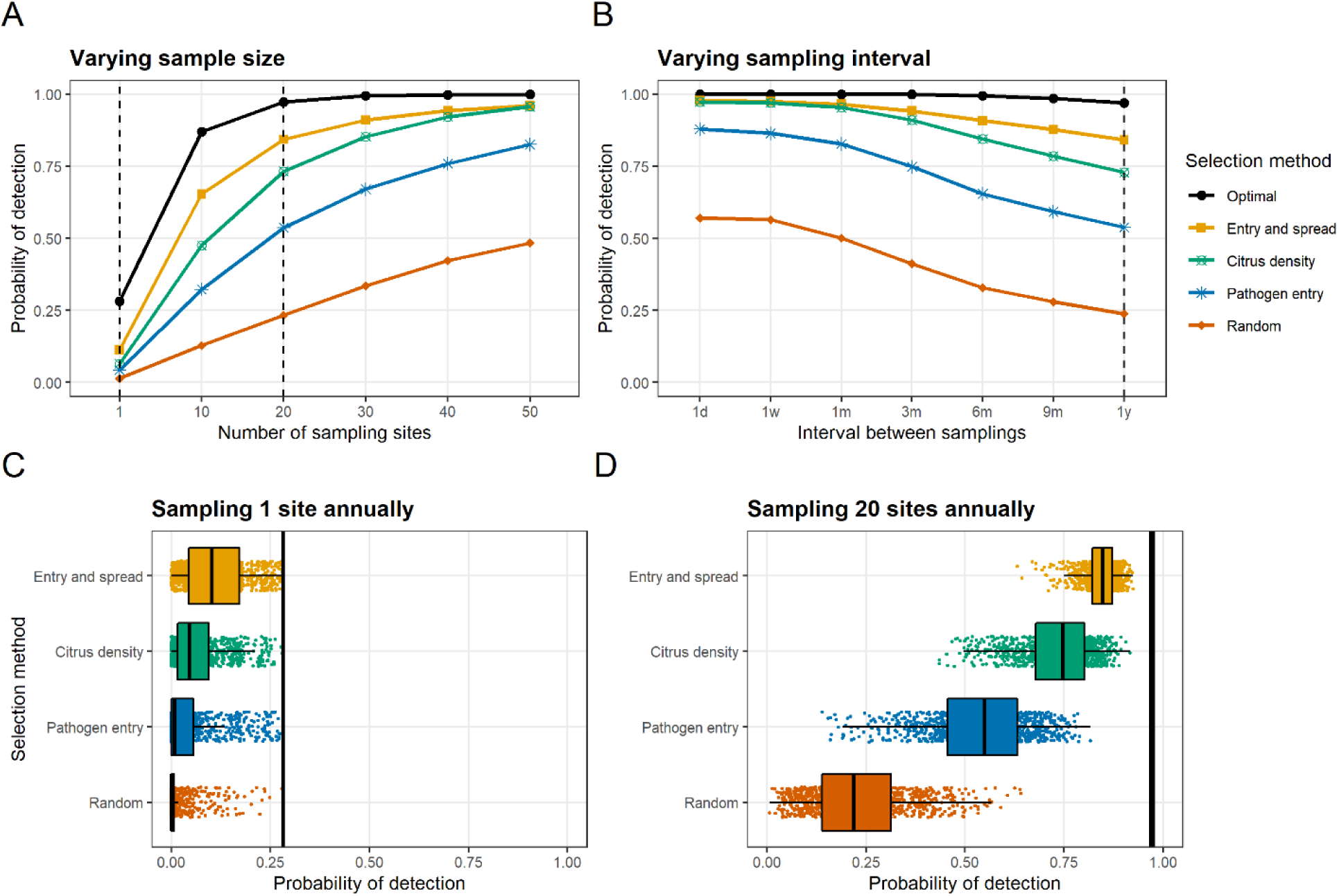
Impact of varying sample size and sample frequency on detection probability. These plots show the detection probability when sampling parameters are adjusted, for both the optimised approach and a selection of different conventional targeted approaches using a variety of different metrics. Plots A and B show the mean probability of detection for a variety of different sampling approaches when the total number of sampling sites (A) or the interval between consecutive sampling rounds (B) is varied. All non-optimised estimates in plots A and B are the mean of 1,000 sampling realisations where the probability of site selection was based upon the site-specific measure of interest. These measures are the product of travel census probabilities and citrus density (‘Entry and spread’); citrus density; probability of entry according to the travel census model (“Pathogen entry”), or random selection from the landscape. The optimised estimates are the mean of 10 optimisation runs. The dashed lines in Plots A and B indicate the situations assessed in Plots C and D, which show the variation in the individual estimates of the detection probability when a single site (left) or twenty sites (right) are visited each year. The ‘optimal’ probabilities of detection under each of these conditions over 10 optimisation runs are shown as (overlaid) solid vertical lines.

**Figure 4.**
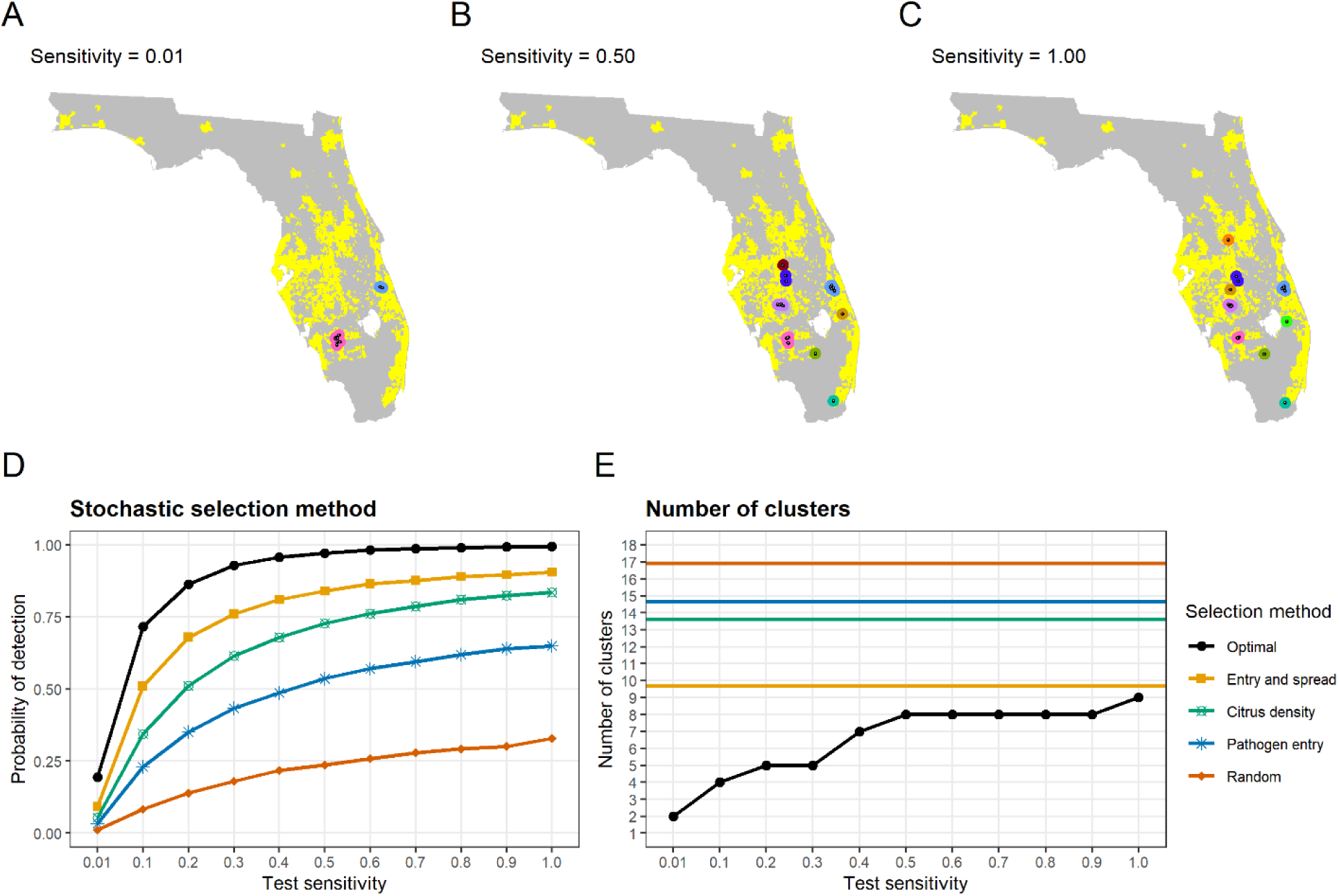
Impact of varying test sensitivity on detection probability and sampling pattern. These plots show the distribution of the optimised strategy for low, moderate, and high detection method sensitivities. They also show the detection probability and number of clusters for a single optimisation run and a variety of alternative targeted strategies using different risk metrics. Plots A, B, and C show the optimal distribution of sampling sites and clusters (points within 20km of each other) when the probability of correctly identifying any sampled infected tree (the diagnostic sensitivity) is low (0.01; plot A), medium (0.50; plot B), and high (1.00; plot C). Plots D and E show the impact of varying the test sensitivity on the overall mean probability of detection (C), and the total number of clusters (D), for both the optimised sites and sites selected using different risk metrics. These metrics are the product of travel census probabilities and citrus density (‘Entry and spread’); citrus density; probability of entry according to the travel census model (“Pathogen entry”), and random selection from the landscape. Due to stochasticity in the risk metric estimates, the horizontal lines in plot D show the mean number of clusters over 1,000 sampling realisations. The optimised results correspond to a single optimisation.

**Figure 5.**
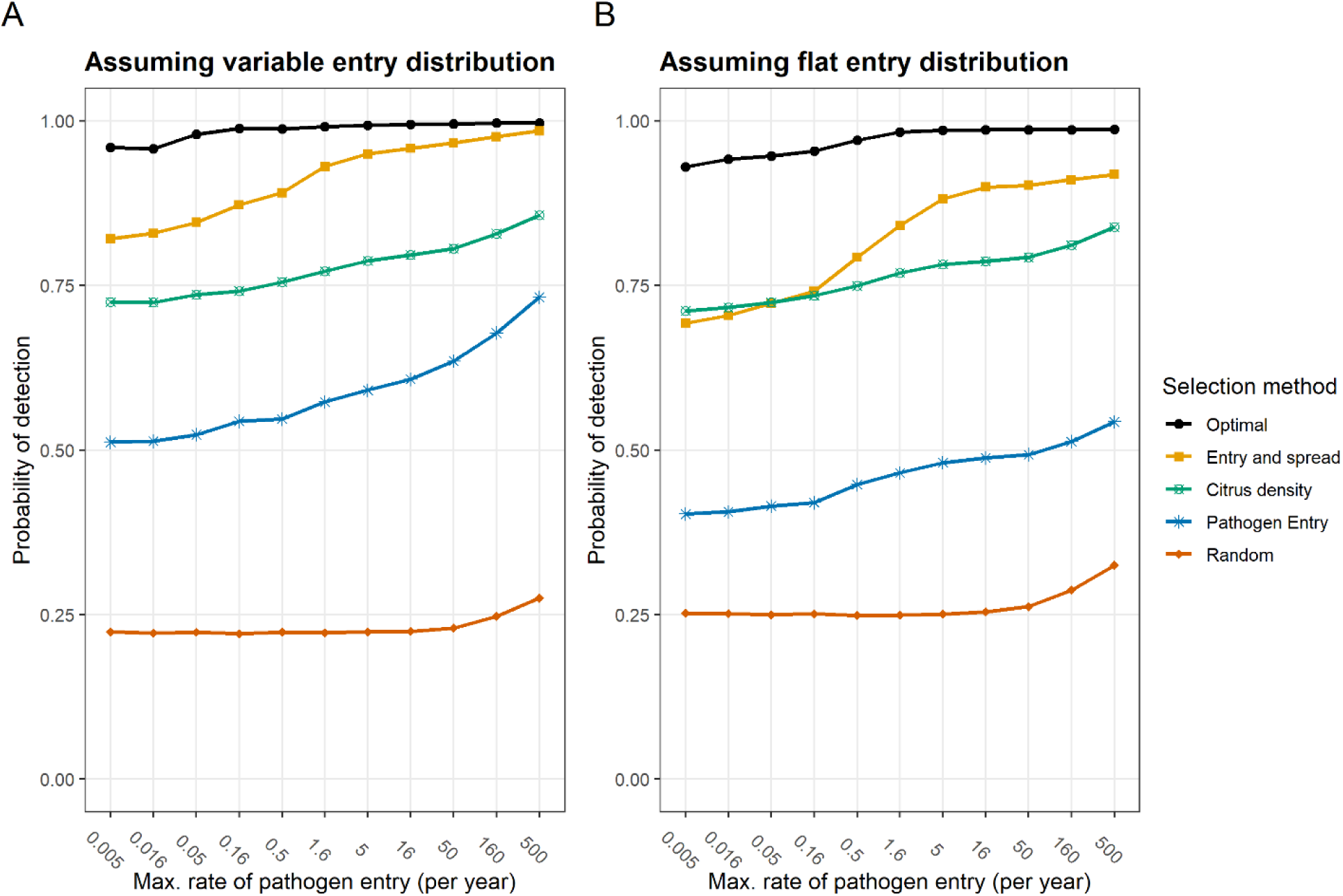
Impact of incorrect assumptions on performance of different methods. These plots show the mean detection probability under different site selection methods for different true rates and patterns of pathogen entry. For each selection method, sites were selected under the baseline model assumptions (i.e. a maximal rate of pathogen entry equal to 0.05/year and a distribution of entry based upon the Travel Census model). The mean of 100 selections are shown for all methods except the ‘optimal’ estimate, which represents the output of a single optimisation run. The risk metrics used for conventional targeted selection represent the product of travel census probabilities and citrus density (‘Entry and spread’); citrus density; the probability of entry according to the travel census model (“Pathogen entry”), and random selection from the landscape. Plot A shows the impact of different rates of pathogen entry given that the distribution pattern was correct, and Plot B shows the same given that the assumed distribution pattern was incorrect (and entry was only affected by citrus density). In all cases, although higher rates result in a higher detection probability in general, the optimised approach (even when based on an incorrectly specified model) outperforms all alternative methods for all entry rates considered.

### Epidemiological simulations

Our model simulates the early stages of an epidemic: up to a pre-defined ‘threshold prevalence’ of 1%, which was selected to indicate a point at which control would no longer be possible. Since the model is stochastic, we simulate not only the spatial spread within a landscape over time, but how this varies from one epidemic to the next. Taking the average over all of these epidemics, we find that the distribution of entry sites (either throughout the citrus landscape, or focused on particular ‘high risk’ areas) affected which sites were more or less likely to be infected by the end of the simulation (Figure 1 and Supplementary Figure 1).

### Comparison with conventional targeted surveillance

To compare our method with more conventional targeted surveillance strategies, we used a variety of methods to quantify the infection risk for each site, and selected sites for surveillance based on these ‘risk metrics’. We considered four risk metrics, each representing a different level of knowledge of the likely presence of the pathogen in each site:

a. Random sampling throughout the simulated landscape (indicating no knowledge of risk factors for infection).
b. Relative rate of pathogen incursion, excluding citrus density (indicating known areas of likely pathogen entry).
c. Citrus density (indicating known areas of likely pathogen establishment and spread).
d. Product of citrus density and relative rate of incursion (indicating known areas of likely pathogen entry, establishment, and spread).

As a result of spatial autocorrelation (i.e. the fact that the risk status of any given site is likely to be similar to that of surrounding sites), we found that simply selecting the sites with the highest risk tended to result in a small number of clusters of sites (Figure 2). To avoid this, we therefore instead used a selection method where the probability of site selection was proportional to the risk metric, as has been used previously for targeted surveillance^20^. This resulted in a greater spread of selected sites (i.e. a larger number of distinct ‘clusters’ of sites) (Figures 2 and 4), and generally gave a higher detection probability than the ranking approach (full results not shown). Although the detection probability of optimised sites was consistently higher than that for sites selected using more conventional methods, the risk metric calculated as the product of citrus density and relative rate of incursion gave the closest detection probabilities to the optimised approach. As expected, random sampling gave the lowest mean detection probability – demonstrating the value of targeted surveillance strategies for early detection surveillance.

### Varying surveillance characteristics

Under the baseline model (a low rate of pathogen entry, focused on high risk sites, with 20 sites surveyed annually using a detection method with a sensitivity of 0.5), the optimised mean detection probability approached 1 (Figure 4). However, increasing the number of sites visited, reducing the sampling interval, or increasing the probability of correctly identifying infected hosts (i.e. the diagnostic sensitivity of the detection method) all increased the mean probability of detection under the optimised strategy. We also found that when the diagnostic sensitivity was high, surveys should be spread throughout the citrus landscape. However, as this was reduced, surveys become increasingly concentrated in a small number of ‘hot spots’, where higher rates of pathogen entry intersected with higher citrus density (Figure 4).

### Varying the rate and distribution of pathogen entry

Increasing the rate of pathogen entry into the state decreased the number of clusters of surveillance sites (Supplementary Figure 3). This effect was more pronounced when ‘high risk’ entry sites were present, reflecting the end prevalence estimates in these sites from the model simulations (Supplementary Figure 1). We identified three consistent clusters of surveillance locations in citrus growing regions to the northeast, northwest, and southwest of Lake Okeechobee, as well as clusters to the east of the lake and in the centre of the peninsula. The detection probability increased slightly as the rate of pathogen entry increased, and was higher when the distribution of introduction points was variable than when it was flat (Supplementary Figure 2), likely reflecting the higher site-specific prevalences (and therefore higher achievable detection probabilities) in these situations. We found that mis-specifying the rate of pathogen entry (i.e. assuming that the baseline model was correct) had a relatively small impact on the performance of the optimised method, which consistently outperformed all other selection strategies (Figure 5). In all cases except random sampling, higher rates of introduction were associated with higher detection probabilities even when site selection was based upon an incorrect model (Figure 5).

## Discussion

### Model-informed surveillance

In the current report, we describe a novel method of identifying how best to deploy surveillance efforts in order to detect epidemics of exotic pathogens at an early stage. We also explore how certain epidemiological and surveillance characteristics impact on the optimal surveillance strategies. Our method links the output of an epidemiologically-informed spatially explicit simulation model capable of reproducing early stage pathogen spread with an optimisation routine informed by specified surveillance parameters^33^. Our key finding is that it is generally best to avoid ‘putting all your eggs in one basket’ when planning surveillance, and that surveillance resources should generally be spread throughout the landscape to cover all areas of risk (Figure 2). This is an important message since many surveillance programs in plant health are typically disproportionately targeted to a small number of high-risk areas, such as areas immediately adjacent to current outbreaks or surrounding ports of entry. Sub-optimal deployment of surveillance resources such as this can be ill-afforded at a time when the number of plant pests and pathogen threats is rising.

Our method also goes beyond the simple maxim of not putting all eggs in one basket, by suggesting precisely which number of baskets should be used and where they should be. We find that the answer to these questions is particularly affected by the performance of the detection method being used (Figure 4), whereas the rate and distribution of pathogen entry is less important (Figure 5, Supplementary Figure 2). Although the exact selected sites varied slightly when the optimisation algorithm was run repeatedly on the same model output, the general locations and arrangement were very similar, and resulted in no perceivable change in the final detection probability (Figure 3) – indicating that the optimisation was robust to slight differences in site selection. We also found that that there was relatively little impact of making the wrong assumption regarding the rate and distribution of pathogen entry on the detection probability of the optimised strategy (Figure 5). Interestingly, we also found that the detection probability for all site selection methods increased as the rate of pathogen entry increased (Figure 5) – likely reflecting a greater spread of infected sites due to relatively less spread within the state.

### Optimisation outperforms conventional targeted surveillance strategies because it considers the system as a whole

For all scenarios assessed (including those based on an incorrectly parameterised model), the optimised surveillance strategy outperformed all strategies using risk metrics (Figures 2, 3, 4, and 5). This results from the optimisation being able to explicitly capture relevant aspects of the transmission and detection process and evaluate the surveillance plan whilst considering the system as a whole. Under more conventional targeted surveillance strategies, we found that the best risk metric to use to select sites was calculated as the product of the relative probability of introduction and the citrus density (which effectively describes the expected relative rate of pathogen entry in the model) (Figures 3 and 4). This metric also gave a clear improvement in comparison to random sampling or selection based upon relative rate of pathogen entry alone – reiterating the importance of considering pathogen epidemiology (both introduction and onwards spread) when selecting surveillance sites. This approach shares characteristics with our previous targeted surveillance strategy of quantifying risk as the product of the introduction probability and the magnitude of onward spread if introduction occurs^20^.

### Uncertainty in the rate and pattern of introduction has a relatively small impact on where best to conduct surveillance

Although the ability to explicitly account for the processes of pathogen entry (‘primary infection’) and onwards spread (‘secondary infection’) is a particular strength of our method, in some cases there may be considerable uncertainty and/or variability in these processes, making them difficult to parameterise. We therefore investigated how the rate and distribution of pathogen entry impact on the optimal surveillance strategy. Although the impact of changing these parameters was relatively low, we found that the optimal surveillance sites tended to be clustered in the sites of highest citrus density for high rates of pathogen entry, with surveillance in lower citrus density areas only being promoted as the rate was decreased (Supplementary Figure 2). Despite these differences, the impact of mis-specifying the rate and distribution of pathogen entry had a relatively small impact on the overall detection probability, which remained high in all scenarios considered (Figure 5). Investigation of the impact of changes in secondary spread patterns, as well as spread within and between different groups of hosts, will be considered in future work.

### The performance of the detection method should be considered when planning where to conduct surveillance

In contrast to the limited impact of the rate and pattern of pathogen entry, our analysis shows that the performance of the detection method (i.e. the diagnostic sensitivity) has a considerable impact on the optimal arrangement of surveillance resources, as well as on the overall detection probability. This finding is due to spatial autocorrelation in the status of individual sites, meaning that the status of each individual surveillance site is not independent of that of nearby sites. Put simply, each site inspected provides some information on the status of nearby sites, but the amount of this information decreases as the diagnostic sensitivity of the detection method and/or the number of samples taken per site decreases (and the probability of detecting infected hosts decreases). This means that the optimal surveillance strategy for any given pathogen will differ when a highly sensitive detection method is used, compared to when the sensitivity is low – with focused sampling in a smaller number of clusters of sites advisable in the latter case. In cases where the test sensitivity and the number of hosts inspected taken per site are low, the value of optimising the surveillance strategy is reduced, and conventional targeted surveillance strategies may be more acceptable (Figure 4). Further work will investigate the impact of detection lag periods^15,34^ on the optimal deployment of surveillance resources.

### Conclusions for HLB surveillance

Under the baseline assumptions regarding spread and detection, our method suggests that the highest probability of early detection of HLB prior to 2005 would have been achieved by focusing surveillance efforts in eight spatial clusters, mainly located in regions of high commercial citrus density in the centre of the peninsula (Figure 2). We estimate that this arrangement of surveillance sites would have given a high chance (97%) of detecting incursions before a statewide prevalence of 1% was reached. This suggested arrangement differs from the actual surveillance implemented prior to the first detection of HLB in Florida in 2005, which was focused to the southeast of Lake Okeechobee, around Tampa in the west of the peninsula, and around Orlando^32^ - none of which are suggested by our method. However, we note that the site of the first detection^32^ was close to the southernmost of our predicted sites (Figure 2), and subsequent positive detections to the northeast and southwest of Lake Okeechobee^35^ were also identified as surveillance targets using our method (Figure 2).

### Surveillance aims

We focus our attention in the current report on surveillance for early detection, rather than other surveillance aims such as prevalence estimation or spatial delimitation^36^ (although our method can be adapted to a variety of surveillance aims, which will be explored in future work). This allows us to concentrate on early stage pathogen spread, when spread dynamics are more predictable and thus easier to model (although we appreciate that there may be uncertainty in parameter estimates in these stages^37^). It also allows us to better explore the impact of epidemiological and diagnostic parameters on the optimal deployment of surveillance resources for early detection (regardless of pathogen impact or required control measures). By considering pathogen spread up to a fixed ‘maximum acceptable prevalence’, our method fits in well with the maximum prevalence thresholds commonly specified when planning conventional regulatory surveillance for regulated pathogens^38,39^. Although optimisation routines have been applied to simulation models in order to improve early detection surveillance for invasive species in other studies^26–30,40^, most of these have focused on estimating the optimal balance of early detection surveillance and control intensity required to minimise the total economic impact of the invasive species. As well as not being able to consider optimal surveillance in isolation of control strategies, these approaches are generally less able to explicitly model pest introduction and spread in a realistic landscape, which limits their use in precise targeting of surveillance activities. Although some recent studies have considered the spatial characteristics of optimised surveillance in more detail, these have either been applied to simulated landscapes^41–43^ or have been based on network modelling approaches^44,45^ which are less able to capture spatial spread patterns. Although the current study is intended to explore optimal surveillance deployment rather than make concrete suggestions for implementation, further work will apply our method to current ongoing threats, such as the spread of Las in California^8^ and *X. fastidiosa* in Italy^10^.

## Methods

### Summary

We consider here how best to deploy surveillance resources in order to maximise the probability of detection before a specified ‘maximum acceptable prevalence’ is reached. To do this, we developed a grid-based stochastic spatially explicit landscape-scale model and repeatedly simulated pathogen spread through this landscape until this prevalence was reached. Although the model is pathogen-generic, we parameterised it to replicate early stage spread of *Candidatus* Liberibacter asiaticus (Las) in Florida. We then used an optimisation routine (simulated annealing) in order to identify which arrangement of a specified number of sites would give the highest mean probability of detection over all simulation model realisations, given a particular frequency of sampling using a detection method with known performance characteristics (the probability of correctly identifying infected hosts (the diagnostic sensitivity) and growth in detectability over time).

### Simulation Model

The simulation model is described in more detail in the Supplementary Information. The simulation model runs on a gridded landscape of 1km x 1km cells, each containing a density of citrus informed by maps of commercial citrus densities (provided by the US Animal and Plant Health Inspection Service) and estimates of residential citrus densities (calculated from population data at the census tract level obtained from the US Census Bureau^46^). Individual cells transition stochastically from a susceptible to an infected status in continuous time, driven by pathogen spread from outside the landscape as well as secondary spread within the landscape. Secondary (between-cell) spread occurs according to an exponential dispersal kernel, fitted to data as described in the Supplementary Information. Following first infection, each cell becomes more infectious (and detectable) as infection bulks-up locally, again at a rate parameterised using available data. This increase in infectiousness and detectability following detection is deterministic, allowing us to estimate the proportion of both infected and symptomatic plants in each cell at any point in time (given the timing of infection in each cell is known, which will vary for each simulation run). By running this stochastic model a large number of times, we are able to capture the inherent variability in spatiotemporal spread and explicitly account for this when considering the optimal arrangement of surveillance sites.

### Calculating the probability to detect at least one case of the epidemic

We used outputs from the simulation model to estimate the probability of detection before the statewide prevalence threshold was reached, *p*(Ω, *n*, Δ*t*), given a sampling pattern consisting of a set Ω of *N* locations from within each of which *n* samples are taken every Δ*t* units of time.

We assume the proportion of detectable hosts in any site increases logistically following first infection, meaning – in site *L* at time *t* in model iteration *i* – the proportion of detectable hosts is given by

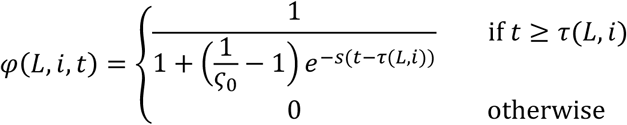

in which *τ*(*L*, *i*) is the iteration-specific time at which site *L* first becomes infected, *ς*_0_ is the proportion of detectable sites at the time of first infection, and *s* is the rate of increase in detectability.

This allows us to quantify the probability of failing to detect infection in site *L* at time *t* as

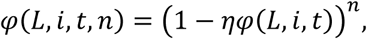

in which each of the *n* samples has a fixed probability of correctly identifying a detectable host (*η*).

The probability of not detecting across the entire sampling pattern is therefore given by

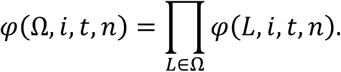

The probability of not detecting in iteration *i* is therefore given by

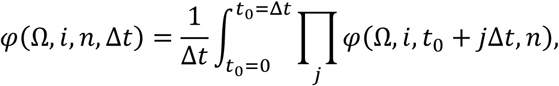

in which the averaging done by the outer integral accounts for uncertainty in the start of sampling (*t*_0_) relative to the time of first introduction of disease anywhere in the landscape (*t*=0), and the inner product runs over all values of *j* until the simulation-specific prevalence threshold has been exceeded.

In simulation *i* the probability of detecting disease given a particular spatial arrangement (Ω), timing (Δ*t*) and local intensity of sampling (*n*) is the complement of this probability, i.e.

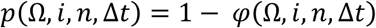

Our final estimate of the effectiveness of any sampling pattern can therefore be obtained by averaging over the *M* simulation runs we consider, i.e.

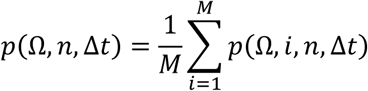

This was used as the objective function in the optimisation algorithm, which therefore identifies the components of Ω.

### Optimisation algorithm

We considered the optimal arrangement of surveillance sites as the selection of 1km square grid cells which gave the highest estimate of *p*(Ω, *n*, Δ*t*), given the sampling intensity (number of sites (*N*), number of samples per site (*n*), and the frequency of sampling (Δ*t*)) and the performance of the detection method. We allowed each site to be selected only once. Because complete enumeration of all possible arrangements is not feasible for a problem of this scale, we used a stochastic optimization algorithm, simulated annealing, with an exponential cooling schedule^47^ to approximate the best arrangement of sites. More details of the algorithm are provided in the Supplementary Information.

### Application to HLB

Parameter values for the simulation model (Table 1) were selected using a combination of statistical model fitting, iterative parameter adjustment, and published data - and are described in more detail in the Supplementary Information. Because of the level of uncertainty in the rate and distribution of pathogen entry into the state (especially resulting from informal, unreported, host movements), we ran different scenarios for these parameters. We considered ‘low’, ‘moderate’ and ‘high’ rates of pathogen entry – for each of which, we modelled two different spatial patterns of entry:

i. A fixed rate, meaning that citrus density alone determined the relative rate of pathogen entry.
ii. Variation in rate according to a probabilistic model of likely entry sites (the ‘travel census model’). This model describes the relative risk of introduction of Las into each census tract of Florida by accounting for the movement of people into the state from other parts of the world where HLB is endemic^23^. The model predictions were based on data from 2010.

**Table 1.** Description of parameters used in the model.

| Interpretation | Value | Rationale |
| --- | --- | --- |
| **Simulation model** |  |  |
| Proportion of citrus in cell. | [Spatial grid] | Estimated as the sum of the commercial and residential citrus density. |
| Citrus density threshold for spread simulation. | 0.0025km<sup>2</sup> | Approximately 1/16<sup>th</sup> of a typical Floridian citrus grove. Also, this density is around the suspected minimum for citrus canker spread in residential trees in Miami<sup>49</sup>. |
| Maximal overall rate of pathogen entry into the state. | 0.05/year (around one entry every 200 years)<br>0.50/year (around one entry every 20 years)<br>5.00/year (around one entry every two years) | These rates were selected in order to represent a variety of different situations. The inverse of each rate relates to the mean number of entries per year if all cells were full of susceptible hosts. Estimates of the modelled rate based on the mean citrus density of 0.1 are shown in parentheses. |
| Initial infectivity following cell infection. | 0.006 | Fitted to data on the spread of disease in Devil's Garden, assuming a logistic increase in infectivity. See Supplementary Information. |
| Relative rate of pathogen entry. | [Spatial grid] |  |
| Rate of increase in infectivity per cell. | 1.25 infections / infected / year | Fitted to data from Devil's Garden, assuming a logistic increase in infectivity and a 6-12 month asymptomatic period. See Supplementary Information. |
| Secondary spread parameter (can be interpreted as the maximal number of cells infected by a single infected cell). | 1,000/year | Selected by fixing all other parameters and adjusting until the model simulated statewide spread in 10 years (see Supplementary Information). |
| Mean pathogen dispersal distance (assuming exponential kernel). | 20km (in two dimensions) | Estimate taken from recent study of pathogen spread<sup>20</sup>. |
| Maximum prevalence for simulation model. | 1% | Selected as representative of a relatively low prevalence which is also able to capture the impact of spatial autocorrelation following introduction. |
| Number of epidemic realisations to run. | 1000 |  |
| **Detection model** |  |  |
| Initial probability of detection. | 0.006 | Fitted to data from Devil's Garden, assuming a logistic increase in detectability. See Supplementary Information. |
| Rate of increase in probability of detection. | 1.00 detectable / detectable / year | Fitted to data from Devil's Garden, assuming a logistic increase in detectability. See Supplementary Information |
| Number of sites visited each sampling round. | 1<br>10<br>20<br>30<br>40<br>50 |  |
| Number of samples collected per site each sampling round. | 50 | Considered a reasonable intensity of surveillance within a 1km<sup>2</sup> area. |
| Probability of detection if infected host sampled. | 0.5 | Assuming visual inspection. Estimate obtained from a comparison of PCR and visual inspection for detection of infection in Floridian citrus groves<sup>51</sup>. |
| Interval between sampling rounds. | 1/365 years<br>7/365 years<br>1/12 years<br>0.25 years<br>0.50 years<br>0.75 years<br>1 year | Daily<br>Weekly<br>Monthly<br>Quarterly<br>Biannually<br>Every nine months<br>Annually |
| **Optimisation** |  |  |
| Initial temperature. | 10 | Identified by adjusting parameter estimates and inspecting the progression of the objective function (Supplementary Figure 3). |
| Rate of cooling. | 0.999 |  |
| Number of iterations of simulated annealing algorithm. | 100,000 |  |

When parameterising the detection process, we considered a ‘baseline’ scenario in which detection was through visual inspection (as is standard for most plant pathogens^48^) of 50 trees in 20 sites annually (i.e. 1,000 trees in total per year), as shown in Table 1. The optimisation algorithm itself was parameterised by applying this baseline detection model to a simulation model parameterised using the values in Table 1 with a low pathogen entry rate, distributed according to the travel census model. We then varied the initial temperature and the cooling parameters of the algorithm and ran the optimisation for 100,000 iterations and inspected the trace of the detection probability. We found little impact of varying these parameters on the objective function of the optimised solution, although the trace plots differed (see Supplementary Figure 3). The optimisation algorithm parameters used are shown in Table 1.

### How is the optimal surveillance strategy influenced by risk, detection methods, and epidemiology?

In order to identify the impact of epidemiological and surveillance system characteristics on site selection, we used the following two methods to characterise the optimal sites:

- Estimating the mean detection probability. This was performed as described above, using a separate dataset of 1,000 iterations to that which the optimisation was applied.
- Visualising and quantifying the spatial arrangement of selected sites by identifying ‘clusters’ of selected sites. We used a single-linkage agglomerative clustering method to group sites within 20km of each other (this distance was selected as representative of the mean annual dispersal distance of HLB in Florida^20^). The distribution of these clusters assists in the visualisation of the general arrangement of selected sites, and the total number of clusters provides a useful summary statistic (with lower values indicating more clustering).

We evaluated how well more conventional targeted surveillance approaches perform in comparison to the optimal sites by creating ‘risk metrics’ accounting for a range of different levels of knowledge about the entry, establishment, and spread of Las (described in the Results). These metrics were allocated to individual sites and the specified number of sites selected (without replacement) with probability proportional to the metric. We repeated this process 1,000 times for each metric and recorded the detection probability for each run using the dataset of 1,000 runs used to estimate the detection probability for the optimal sites.

## Acknowledgements

S.R.P. was part-funded by the USDA APHIS farm bill grant (17-8130-0570-CA) and Defra. F.v.d.B. received funding from the BBSRC whilst at Rothamsted Research.

