## Supplemental Information for "Optimising risk-based surveillance for early detection of invasive plant pathogens"

Supplementary Information: Methods

#### Model formulation

The model is a spatially-explicit susceptible-infected model, which tracks – in continuous time – the infection status of each 1km x 1km cell. The model accounts for heterogeneous densities of citrus and rates of pathogen entry, as well as local bulking-up of pathogen densities at any location and dispersal between cells.

The model runs on a rectangular grid of 1km$^2$ cells, of extent 672 x 732, to bound the whole of mainland Florida (i.e. excluding the Florida Keys) – encompassing a total of 148,089 cells. The model takes two spatially-heterogeneous inputs, which specify – for each location $L$ – the following (fixed) quantities:

- $\rho_L$= proportion of the cell $L$ containing citrus;
- $\varepsilon_L$= rate of pathogen entry on the cell $L$.

Only the 35,157 cells across the landscape with $\rho_L \geq 0.0025$ (i.e. which contain at least 2,500m$^2$ of citrus) are tracked. This set of active cells is denoted as $A$. Filtering out cells containing only very small densities of host plants led to a significant increase in computational speed with minimal impact on the spatiotemporal pattern of simulated pathogen spread, due to the low probability of infection in these cells even if included. Although this feature of the model fragments the landscape, spread within the main commercial citrus growing area in the centre of the Floridian peninsula remains well connected, and long-range spread is still captured through repeated pathogen incursions.

Cells are partitioned into two disjoint sets: infected $(I)$ and susceptible $(S)$, with $I \cup S = A$. We assume the first infection of a cell leads to $\sigma_0 = 0.006$ of the citrus within it becoming infected, and that thereafter the pathogen bulks-up logistically at rate $r$. Following first infection, within-cell spread of the pathogen is assumed to be dominated by local spread, which occurs deterministically[3]. This simplifies many of the calculations and allows large numbers of simulations to be run in a shorter time period - which is a large advantage of our current model. In particular, we assume that if cell $L \in I$ is first infected at $t_L$, at any subsequent time the density of infected host is given by:

$$\omega_L(t) = \frac{\rho_L}{1 + \left(\frac{1}{\sigma_0} - 1\right) e^{-r(t-t_L)}}$$

The only stochastic transition tracked by our model is pathogen spread between cells. The time-dependent net rate at which any uninfected cell $L \in S$ becomes infected, $\lambda_L(t)$, is:

$$\lambda_L(t) = \rho_L \left( \varepsilon_L + \beta \sum_{\bar{L} \in I, \bar{L} \neq L} K(L, \bar{L}) \omega_{\bar{L}}(t) \right)$$

in which $\beta$ scales the secondary infection rate and $K(L, \bar{L})$ is a dispersal kernel linking uninfected cell $L$ with infected cell $\bar{L}$. We use an appropriately normalised two-dimensional exponential kernel, where:

$K(L, \bar{L}) = \frac{1}{2\pi\alpha^2} \exp\left(-d_{L,\bar{L}} I \alpha\right)$

Here, $\alpha$ is a dispersal scale parameter and $d_{L,\bar{L}}$ is the distance between the centres of cells $L$ and $\bar{L}$. The process of infection is simulated according to an appropriate update to Gillespie algorithm[4,5] which accounts for time-inhomogeneous rates.

We modelled the increase in detectability ($\varphi$) in each cell over time using the same approach described for infectiousness above, with a rate parameter of $s$ and an initial detectability of $\varsigma_0$.

$$\varphi_L(t) = \frac{1}{1 + \left(\frac{1}{\varsigma_0} - 1\right) e^{-s(t-t_L)}}$$

We fit the infection rate, $\beta$, the scale of dispersal $\alpha$, the rates of within-cell bulk up and detectability increase, $r$ and $s$, and the initial cell infectiousness and detectability, $\sigma_0$ and $\varsigma_0$, to spread data as described below.

#### Optimisation algorithm

Our algorithm first randomly selects the required number of sites and calculates the mean probability of detection, $p(\Omega, n, \Delta t)$, for the given sampling arrangement $\Omega_j$ (under the given surveillance parameters, $n, \Delta t$), as described in the main text. We used this detection probability as the 'objective function' in the optimisation algorithm, which needs to be maximised. For a prespecified number of iterations $(J)$, the algorithm proceeds by sequentially replacing a single site with another randomly selected one before calculating the objective function again. The arrangement with the new site is then either accepted or rejected before another site is randomly replaced and the process repeated. Each time, the following Metropolis criterion is used to estimate the probability of accepting the new site:

$$P(\Omega_j \rightarrow \Omega_{j+1}) = 1 \text{ if } p(\Omega_{j+1}, n, \Delta t) < p(\Omega_j, n, \Delta t)$$

$$P(\Omega_j \rightarrow \Omega_{j+1}) = \exp\left(\frac{p(\Omega_{j+1}, n, \Delta t) - p(\Omega_j, n, \Delta t)}{temp_j}\right) \text{ if } p(\Omega_{j+1}, n, \Delta t) > p(\Omega_j, n, \Delta t)$$

If the objective function was equal between iterations, then the new arrangement was accepted with a probability of 0.5. The ‘temperature’ of the algorithm $(temp)$ is multiplied by the cooling rate $(alpha)$ at the end of each iteration (i.e. an exponential cooling schedule[6]). The result of this is that in the early stages of the algorithm, $temp$ is high and so is the probability of accepting ‘worse’ arrangements of sampling sites - thereby encouraging a full exploration of the full parameter space, avoiding any local maxima. As the algorithm progresses, $temp$ decreases and it becomes increasingly likely that worse arrangements are rejected (although there initially remains some freedom to explore the parameter space). In the late stages of the algorithm, all arrangements which give a lower probability of detection are rejected, allowing a good approximation of the true optimal arrangement to be found.

### Model Parameterisation

The citrus density was estimated by summing 1km square gridded data on the distribution of residential and commercial citrus in the state as described in the main text. Following introduction, we assume that the prevalence of infection (and therefore, the infectiousness of the cell) grows logistically. We also assume that the detectability of infection increases logistically, although not necessarily at the same rate. To determine the rate of increase in detectability, we fitted a logistic model to data on the progression of visually detectable infection over time, collected in the ‘Devil’s Garden’ plantation in southern Florida[7,8]. This gave a growth rate of about 0.0028 per day, which is about 1.0 per year. Using data on the development of symptoms over time (described below), and assuming a lag of around six months between infection and first expression of symptoms[9], we assumed that the rate of logistic growth in the prevalence was 1.25 per year. The diagnostic sensitivity of visual inspection was assumed to be 0.5 (inferred from a comparison of PCR and visual inspection for detection of infection in Floridian citrus groves[10].

In order to estimate the rate and pattern of secondary spread, we first assumed that the mean distance of spread (in two dimensions) was around 20km, which corresponds to data obtained in a recent study[11]. We then ran ten simulations of Las spread with identical parameters, restricting initial entry to the four counties found to be infected at the time of first detection (Miami-Dade, Broward, Palm Beach and Martin, in the south east of the state)[12], and assuming a low rate of pathogen entry (0.05), following the distribution predicted by the travel census model. We repeated this process whilst incrementally increasing the rate of secondary spread until the prevalence after ten years exceeded 95%.

### Supplementary Information: Figures

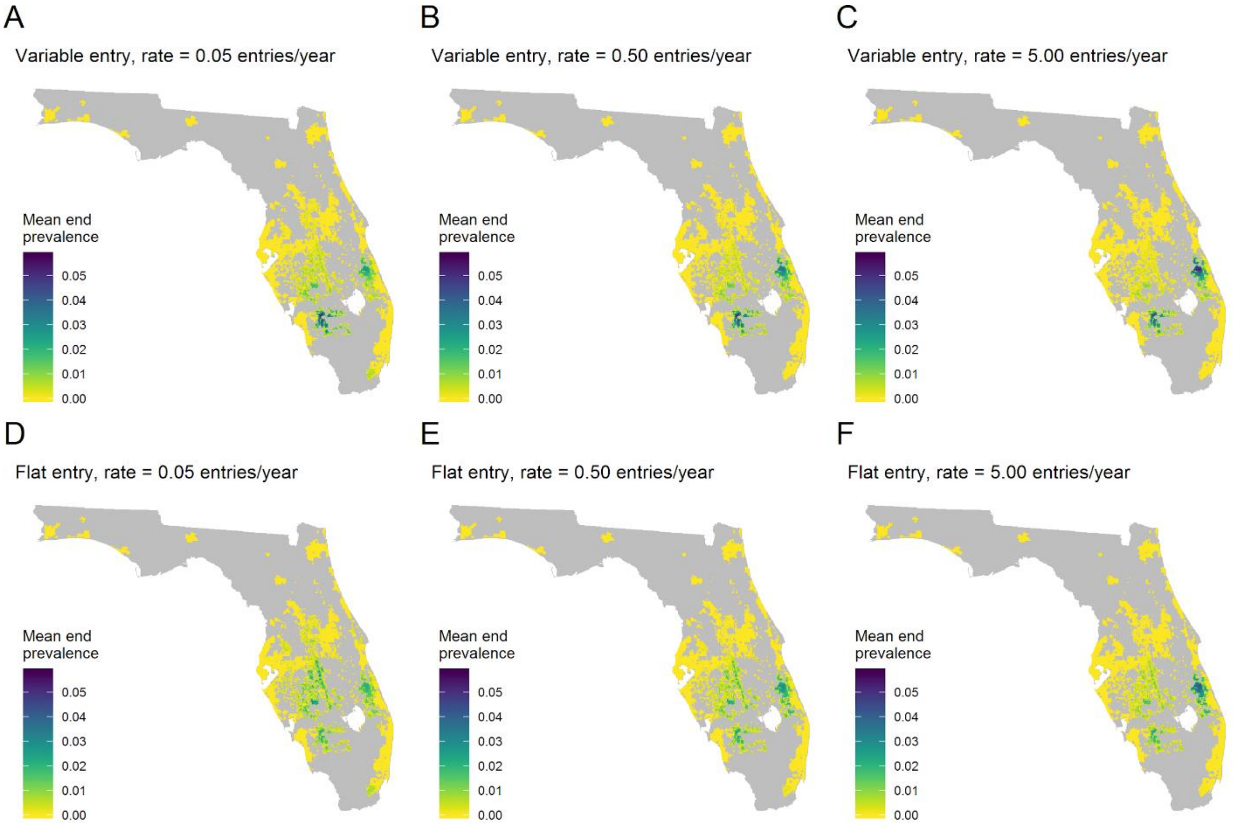


**Supplementary Figure 1. Distribution of mean end prevalence estimates under different patterns of pathogen entry, demonstrating the increase in variability when pathogen entry is more variable.** These plots demonstrate the impact of varying the characteristics of pathogen entry into the State on the mean prevalence at the point the statewide prevalence threshold of 1% is reached. Plots A-C show the mean prevalence when the probability of pathogen entry into any given site is affected by factors beyond just the density of citrus host (in this case, the travel census probabilities), and plots D-F show the distribution when only the citrus density influences the probability of pathogen entry. Plots A and D show a ‘low’ mean rate of pathogen entry up to 0.05 entries per year; B and E show a ‘medium’ mean rate of up to 0.5 entries per year; and C and F show a ‘high’ rate of up to 5 entries per year. Higher rates of entry result in more variability in end prevalence estimates throughout the State.

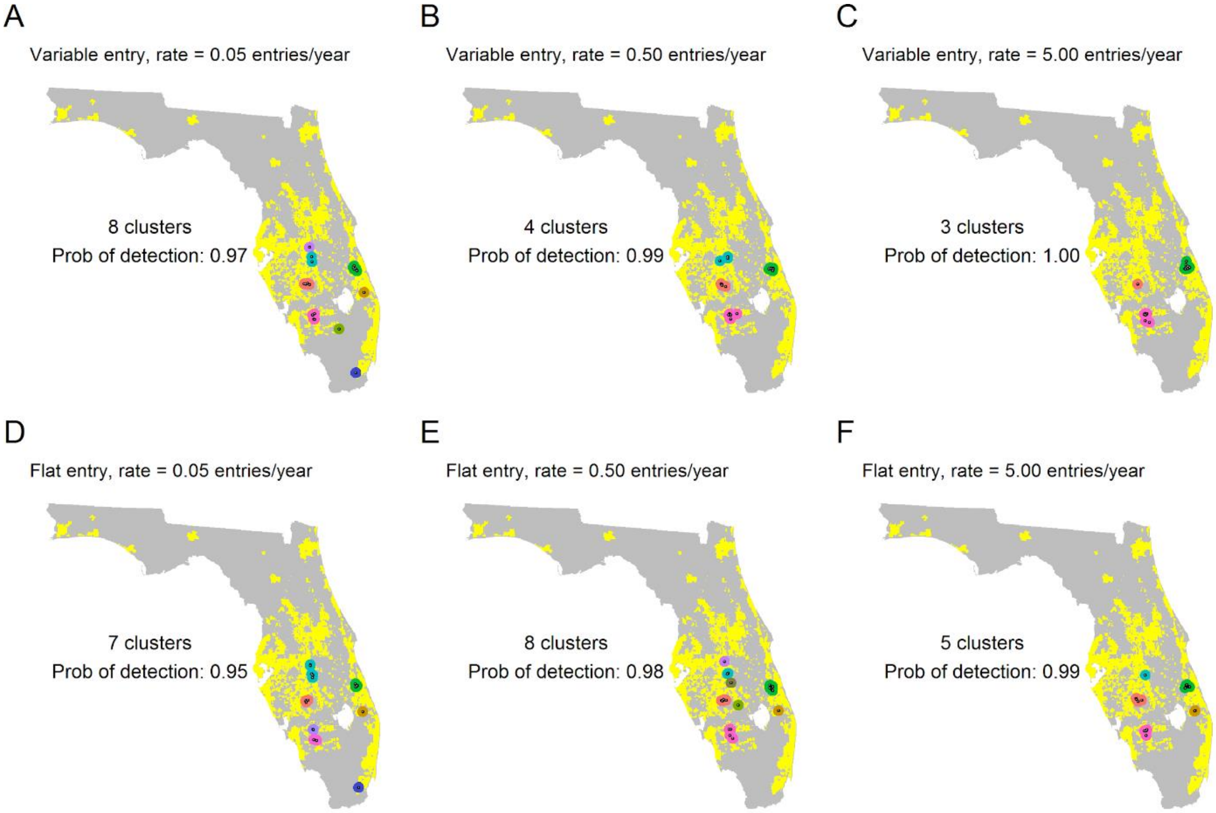


**Supplementary Figure 2. Impact of rate and distribution of pathogen entry on optimal targeting of surveillance.** These plots show the spatial arrangement of optimal sites (taken from a single optimisation run) and clusters of these sites, along with the detection probability, when the rate of pathogen entry is varied. Plots A-C show the distribution when some information on the distribution of introduction sites is available (modelled as the product of travel census probabilities and citrus densities). Plots D-F show the distribution in the absence of this knowledge (assuming establishment is only influenced by citrus density). Plots A and D show a 'low' mean rate of pathogen entry up to 0.05 entries per year; B and E show a 'medium' mean rate of up to 0.5 entries per year; and C and F show a 'high' rate of up to 5 entries per year. Estimates of the number of clusters and the probability of detection under the different sampling patterns are also shown.

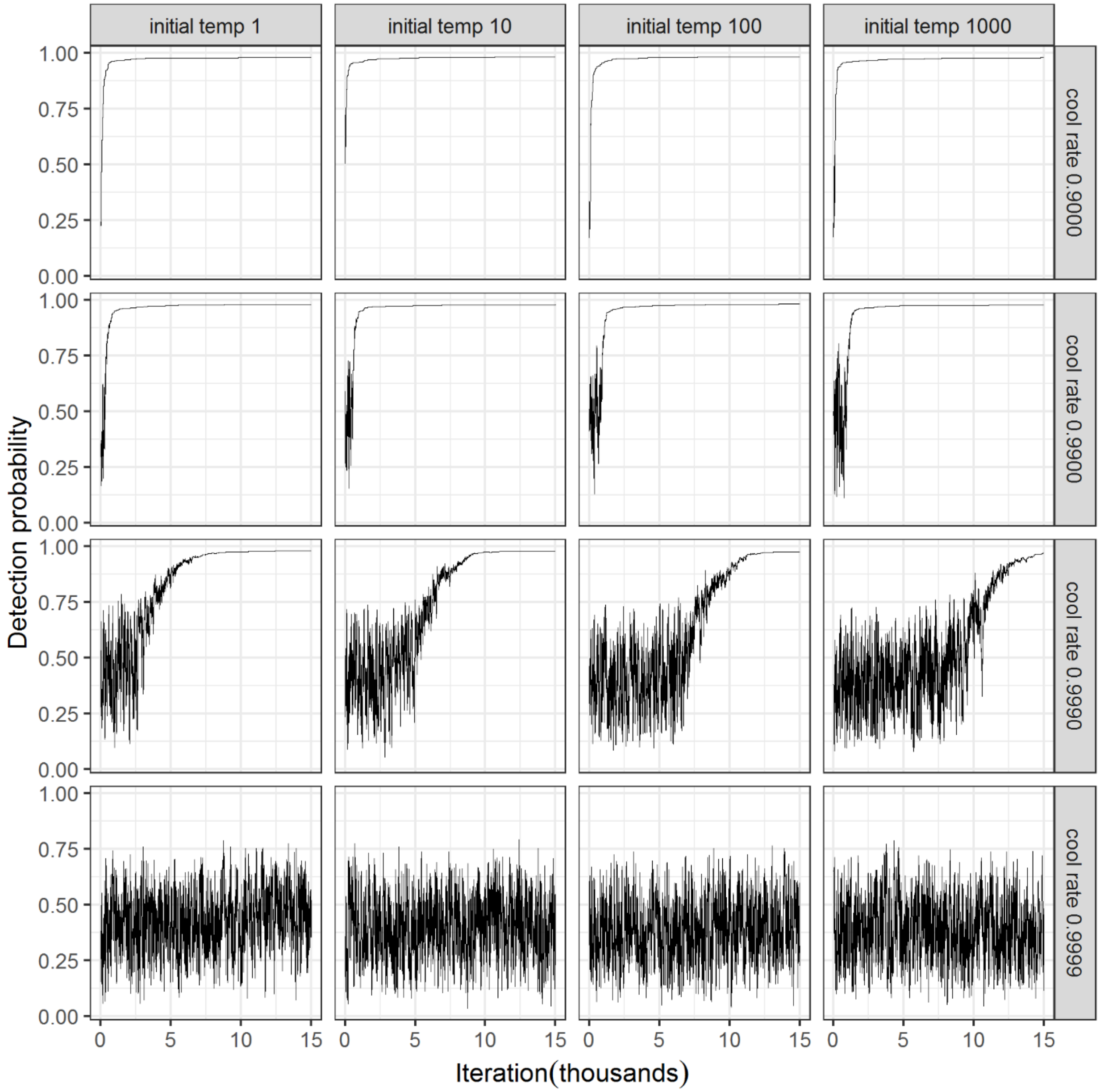


**Supplementary Figure 3. Plots of the 'trace' of the simulated annealing algorithm for a range of different parameter values.** This plot shows the change the detection probability as the simulated annealing algorithm progresses over the first 15,000 iterations, using the baseline simulation and detection parameters. The final selected combination of initial temperature and cooling parameters were 10 and 0.9990, respectively (plot in the second column of the third row)
